# Single-molecule fluorescence-based approach reveals novel mechanistic insights into small heat shock protein chaperone function

**DOI:** 10.1101/2020.02.16.951632

**Authors:** Caitlin L. Johnston, Nicholas R. Marzano, Bishnu Paudel, George Wright, Justin L. P. Benesch, Antoine M. van Oijen, Heath Ecroyd

## Abstract

Small heat shock proteins (sHsps) are a family of ubiquitous intracellular molecular chaperones that are up-regulated under stress conditions and play a vital role in protein homeostasis (proteostasis). It is commonly accepted that these chaperones work by trapping misfolded proteins to prevent their aggregation, however fundamental questions regarding the molecular mechanism by which sHsps interact with misfolded proteins remain unanswered. Traditionally, it has been difficult to study sHsp function due to the dynamic and heterogenous nature of the species formed between sHsps and aggregation-prone proteins. Single-molecule techniques have emerged as a powerful tool to study dynamic protein complexes and we have therefore developed a novel single-molecule fluorescence-based approach to observe the chaperone action of human αB-crystallin (αBc, HSPB5). Using this approach we have, for the first time, determined the stoichiometries of complexes formed between αBc and a model client protein, chloride intracellular channel 1 (CLIC1). By examining the polydispersity and stoichiometries of these complexes over time, and in response to different concentrations of αBc, we have uncovered unique and important insights into a two-step mechanism by which αBc interacts with misfolded client proteins to prevent their aggregation. Understanding this fundamental mechanism of sHsp action is crucial to understanding how these molecular chaperone function to protect the cell from protein misfolding and their overall role in the cellular proteostasis network.

## Introduction

Small heat shock proteins (sHsps) are a diverse and ubiquitously expressed family of intracellular molecular chaperones that play a critical role in the maintenance of protein homeostasis (proteostasis). One of the main roles of sHsps is to bind and trap misfolded proteins to protect cells from irreversible protein aggregation during periods of cellular stress^1–3^. Consequently, sHsp malfunction has been implicated in a number of diseases including cataracts, cancer, motor neuropathies and neurodegeneration^4–6^.

Typically sHsps form oligomeric species in solution and this is thought to be linked to their chaperone function. For example, human αB-crystallin (αBc: HSPB5), an archetypal sHsp and one of the most widely expressed of the 10 human sHsp isoforms, forms large, polydisperse oligomeric ensembles in dynamic equilibrium mediated by subunit exchange^7–9^. These large oligomers are formed from monomeric and/or dimeric building blocks. Many factors, including the presence of client proteins, temperature and post-translational modifications, shift the equilibrium from larger polydisperse oligomers to predominantly smaller oligomers^10,11^.

Whilst it is well established that sHsps can form high-molecular mass complexes with misfolded clients to prevent their aggregation^12–14^, little is known about how these complexes are assembled. It has been postulated that smaller sHsp oligomers have enhanced chaperone activity as a result of their increased exposed hydrophobicity and, therefore, a greater affinity for misfolded and aggregation-prone proteins^15–18^; however, others have suggested that the larger oligomers are chaperone active^19–21^. Thus, it remains unclear precisely how sHsps capture misfolded proteins to form the high-molecular mass sHsp-client complexes observed as a result of their chaperone action.

Studies of monodisperse sHsps from plants (e.g. Hsp18.1 and Hsp16.9), using techniques that include size exclusion chromatography, electron microscopy and native mass spectrometry,have provided important stoichiometric and mechanistic information on the end-stage complexes that these sHsps form with client proteins^22–27^. However, the initial binding events that lead to the formation of these end-stage complexes remain to be resolved and very little is known about the complexes formed between polydisperse mammalian sHsp isoforms and their client proteins. This is primarily due to the heterogeneous nature of these complexes, which may contain a variety of sHsp and misfolded client subunits.

Single-molecule fluorescence techniques overcome some of the difficulties of studying dynamic and heterogeneous systems by facilitating the observation of individual protein-protein interactions. Consequently, such approaches may be advantageous for the study of molecular chaperones^28^, since, in the case of sHsps, they may enable the intial steps of binding with client proteins to be obsereved and therefore the molecular mechanism of chaperone action of sHsps to be revealed. Thus, in this work we have deleveloped and exploited a single-molecule fluoresence-based assay in order to directly observe complexes formed between αBc and a model client protein, the chloride intracellular channel 1 (CLIC1) protein.

Destabilisation of CLIC1, whether through a change in pH or temperature, results in the formation of a folding-intermediate with a high-degree of solvent-exposed hydrophobicity^29,30^, causing it to be decidedly aggregation-prone. This is typical of the client proteins of sHsps that form during times of cellular stress, whereby sHsps bind to these destabilised forms to prevent their aggregation^31^. This led us to exploit CLIC1 as a model client protein for the study of αBc chaperone activity at the single-molecule level. We demonstrate that αBc inhibits the heat-induced amorphous aggregation of CLIC1 and that this inhibitory activity results in the formation of a polydisperse range of αBc-CLIC1 complexes. Employing our single-molecule fluorescence-based assay we have, for the first time, determined the stoichiometries of complexes formed betweenαBc and a client protein, and measured how these complexes change over time. Our results provide evidence for a two-step mechanism of sHsp-client interaction and provide fundamental insight into the molecular mechanisms by which sHsps interact with client proteins to prevent aggregation as part of proteostasis.

## Methods

### Materials, protein expression and purification

All materials in this work were purchased from Sigma Aldrich (St Louis, MO, USA) or Ameresco (Solon, OH, USA) unless otherwise stated. The pET28a bacterial expression vector, containing human αBc wild type (aBc_WT_) or mutant αBc_C176_ were used for expression of the recombinant proteins (Genscript, Piscataway, NJ). The mutant αBc_C176_ was engineered to contain an additional cysteine (compared to aBc_WT_) at the extreme C-terminus to facilitate the site-specific covalent attachment of a fluorescent dye. Plasmids were transformed into competent *Escherichia coli* (*E. coli*) BL21 (DE3) cells. The αBc variants were purified as described previously^32^ and stored at −20°C. CLIC1_C24_ in the pET24a vector was produced via site directed mutagenesis of the wild type genes (Genscript, Piscataway, NJ). The CLIC1_C24_ construct used in this study contained a mutation of one of the native tryptophan residues to phenylalanine (W23F) and mutations of five of the native cysteines to alanines (C59A, C89A, C178A, C191A, C223A); the remaining cysteine (C24) was not modified but used for site-specific fluorescent labelling. The pET24a vector containing CLIC1_C24_ was transformed into *E. coli* BL21 CodonPlus (DE3) RIPL cells and recombinant protein expression was induced by addition of 0.1 mM IPTG and overnight incubation at 18°C. The cells were then harvested by centrifugation at 5000 x g for 10 min at 4°C and the pellet stored at −20°C. Cells were resuspended in 50 mM Tris-base (pH 8.0) containing 100 mM NaCl, 0.5 mg/mL lysozyme and EDTA-free cocktail protease inhibitor, incubated for 20 min at 4°C and then sonicated to further lyse cells and shear DNA. The cell lysate was then clarified by centrifugation twice at 24000x g for 20 min, passed through a 0.45 μM filter and applied to a 5 mL His-Trap Sephadex column (GE Healthcare, USA) equilibrated in 50 mM Tris-base (pH 8.0) containing 5 mM imidazole and 300 mM NaCl. Bound recombinant protein was then eluted with 500 mM imidazole and loaded onto a s75 Superdex size-exclusion column equilibrated in 50 mM phosphate buffer (pH 7.4). Recombinant protein was concentrated, snap-frozen in liquid nitrogen and stored at −20°C until use. The superoxide dismutase 1 (SOD1) used in this work was a gift from Prof. Justin Yerbury (University of Wollongong, Australia).

### In vitro amorphous aggregation assays

*In vitro* aggregation assays were performed to assess the ability of αBc_WT_ and αBc_C176_ to inhibit the amorphous aggregation of CLIC1_C24_. CLIC1_C24_ (30 μM) was incubated in 50 mM phosphate buffer (pH 7.4) supplemented with 10 mM DTT in the presence or absence of varying molar ratios of αBc (1:0.5-1:64, αBc:CLIC1). CLIC1_C24_ incubated in the presence of SOD1 at a 1:0.5 molar ratio (SOD1:CLIC1) acted as a control for the chaperone-specific inhibition of CLIC1_C24_ aggregation. Samples were prepared in duplicate in a Greiner Bio-One 384-well microplate (Greiner Bio One, Freickenhausen, Germany) and sealed to prevent evaporation. The aggregation of CLIC1_C24_ was monitored by measuring the light scatter at 340 nm using a FLUOstar Optima plate reader at 37°C for 20 hr. To quantify the ability of the αBc variants to prevent CLIC1_C24_ aggregation, the percent inhibition of aggregation was calculated using the formula: % inhibition = (Δ*Ic* - Δ*Is*)/ Δ*Ic*) × 100, where Δ*Ic* and Δ*Is* are the change in absorbance in the absence and presence of chaperone at the end of the assay, respectively. The percent inhibition of aggregation afforded by the αBc variants is reported as the mean ± S.D. of three independent experiments.

### Fluorescent labelling of proteins

For single-molecule förster resonance energy transfer (smFRET) experiments, CLIC1_C24_ was labelled with an Alexa Fluor 555 donor maleimide fluorophore (AF555-CLIC1_C24_), and αBc_C176_ was labelled with an Alexa Fluor 647 maleimide acceptor fluorophore (AF647-αBc_C176_). For two-colour single-molecule experiments, CLIC1_C24_ and αBc_C176_ were labelled with Alexa Fluor 647 and Alexa Fluor 488 maleimide fluorophores, respectively. Proteins were fluorescently labelled as previously described with some modifications^33^. Briefly, proteins to be labelled were incubated in 5 mM TCEP and 70% (w/v) ammonium sulphate powder and placed on a rotator at 4°C for 1 hr. Proteins were then centrifuged and the pellet resuspended in degassed buffer A (100 mM Na_2_PO_4_ (pH 7.4), 200 mM NaCl, 1 mM EDTA, 70% (w/v) ammonium sulphate). The protein was centrifuged and the washed pellet was resuspended in buffer B (100 mM Na_2_PO_4_ (pH 7.4), 200 mM NaCl, 1 mM EDTA) containing a 5-fold molar excess of maleimide-conjugated fluorophore. The protein was then incubated on a rotator at room temperature for 3 hr. Following the coupling reaction, excess dye was removed by gel filtration chromatography using a 7 k MWCO Zebra Spin Desalting column equilibrated in 50 mM phosphate buffer (pH 7.4). The concentration and degree of labelling was calculated by UV absorbance or denaturing mass spectrometry (Supplementary Table 1) and stored at −20°C.

### Coverslip preparation and immobilisation of samples for smFRET and two-colour TIRF microscopy

Microfluidic flow cells were constructed by placing PDMS lids on 24 x 24 mm coverslips that had been PEG-biotin-functionalised^34^. Coverslips were functionalised by treatment with 100% ethanol and 5 M KOH, before aminosilanisation was carried out in a 1% (v/v) (3-Aminopropyl) triethoxysilane (Alfa Aesar, UK) solution. PEGylation of coverslips was performed by incubating coverslips with 1:10 mixture of biotinPEG-SVA and mPEG-SVA (Laysan Bio, AL) prepared in 50 mM 3-(N-Morpholino) propanesulfonic acid (MOPS) (pH 7.5) solution for 3 hr. Coverslips were further functionalised by an additional PEGylation overnight before being stored under nitrogen gas at −20°C. Inlets and outlets in the PDMS were prepared using PE-20 tubing (Instech, PA, USA) that allowed washing and addition of samples onto the coverslip surface. Neutravidin (125 μg/ml) was incubated in the flow cell for 10 min, washed with 50 mM phosphate buffer (pH 7.4) supplemented with 6-hydroxy-2,5,7,8-tetramethylchroman-2-carboxylic acid (6 mM, TROLOX) (imaging buffer). To help prevent non-specific interactions of proteins with the coverslip surface, the microfluidic channel was blocked with 2% (v/v) Tween-20 for 20 min^35^ and then washed extensively with imaging buffer. To facilitate immobilisation of His-tagged CLIC1 to the coverslip surface, anti-6X His tag antibody (1 μg/ml) was incubated in the flow cell for 10 min. Finally, pre-formed CLIC1-αBc complexes were diluted 1:1000, incubated in the flow cell for 10 min and washed with imaging buffer to remove unbound protein. To reduce blinking and unavoidable photobleaching of fluorescent dyes during imaging, an oxygen scavenger system (OSS) consisting of protocatechuic acid (PCA, 2.5 mM) and protocatechuate-3,4-dioxygenase (PCD, 50 nM) in imaging buffer was introduced into the flow cell prior to image acquisition.

### smFRET sample preparation, instrument setup and data analysis

To confirm that αBc_C176_ formed complexes with aggregating CLIC1_C24_, smFRET experiments were performed. AF555-CLIC1_C24_ (1 μM) was incubated in the presence of AF647-αBc_C176_ (2 μM) for 20 hr at 37°C. The sample was then diluted 1:1000 in imaging buffer and immediately loaded into a flow cell for TIRF microscopy. Single-molecule measurements were performed at room temperature (approx. 20°C) on a custom-built total internal reflection fluorescence (TIRF) microscope with a Sapphire, green (532 nm) laser that has been previously described^36^. Images were acquired every 200 msec and single-molecule fluorescence intensity time trajectories from multiple fields of view were generated and analysed using a Matlab-based software (MASH-FRET)^37^. Donor leakage into the acceptor channel was corrected during image analysis.

### Single-molecule two-colour sample preparation

Two-colour TIRF microscopy was used to characterise the complexes formed between αBc and CLIC1. To determine how the stoichiometries of αBc-CLIC1 complexes changed over time, 1 μM Alexa Fluor 647-labelled CLIC1_C24_ (AF647-CLIC1_C24_) was incubated in 50 mM phosphate buffer (pH 7.4) at 37°C for 10 hr in the presence of 2 μM Alexa Fluor 488-labelled αBc_C176_ (AF488-αBc_C176_). Aliquots were taken from the reaction at 0, 0.25, 0.5, 0.75, 1, 4, 8 and 10 hr for single-molecule imaging. To examine the effect of chaperone concentration on the stoichiometries of αBc-CLIC1 complexes, AF647-CLIC1_C24_ (1 μM) was incubated under the same conditions as described above except in the presence of varying molar ratios of AF488-αBc_C176_ (0.5:1, 1:1, 2:1 and 4:1 [αBc:CLIC1]) for 8 hr. All samples were diluted 1:1000 into imaging buffer and immediately loaded into flow cells for imaging.

### Two-colour total internal reflection fluorescence microscopy instrument setup and data acquisition

Samples were imaged at room temperature (approx. 20°C) using a custom-built total internal reflection fluorescence microscope system constructed around an inverted optical microscope (IX70, Olympus, Tokyo, Japan). Samples were illuminated by a solid-state 488 nm laser (0.75 W/cm^2^; 150 mW Sapphire 488 nm, Coherent, Santa Clara, CA, USA) and 637 nm laser (6.5 W/cm^2^; 140 mW Vortran, Sacramento, CA, USA), which were aligned and directed off a dichroic mirror (Di01-R405/488/561/635, Semrock, Rochester, NY, USA) to the back-aperture of a 1.49 NA TIRF objective lens (100 x UApoN model, Olympus) mounted on the optical microscope. Fluorescence emission was collected by the same objective and the returning TIRF beam was filtered by a dichroic mirror (Di01-R405/488/561/635, Semrock). Then, incoming emission signals were separated using a dual view of 635 nm cut off dichroic filter (Photometric DV2) that split incoming emission signals into two and directed them to a CCD chip, allowing simultaneous imaging of two colors on each half of the same chip, and passed through appropriate band pass filters (BLP01-488R for AF488 and BLP01-633R for AF647) onto a EM-CCD camera (ImageEM, Hamamatsu, Japan). Control of the hardware was performed using the microscopy platform Micromanager (NIH, USA) and the camera was in frame transfer mode at 5 Hz. Multiple single-molecule movies of each sample were recorded at different fields of view, with images taken every 200 msec. All excitation intensities were kept constant for all samples imaged.

### Two-colour total internal reflection fluorescence microscopy data and statistical analysis

Images were corrected for laser intensity profile and background before intensity time trajectories were generated for all fluorescent molecules using custom-written scripts in Fiji^38^. The initial fluorescence intensity (I_0_) was calculated by averaging the first 50 intensity values (first 10 sec) for all fluorescent proteins identified. Molecules with single photo-bleaching steps for AF647-CLIC1_C24_ and AF488-αBc_C176_ were manually identified and each collectively fit to a Gaussian distribution from which the mean photo-bleaching initial intensity (I_s-mean_) was calculated. The I_s-mean_ values were then used to calculate the number of fluorescently-labelled proteins per point (FPP) using the equation FPP = I_0_/I_s-mean_. At each treatment point (timepoint or concentration) FPP for AF647-CLIC1_C24_ or AF488-αBc_C176_ were combined to determine oligomer size distributions. These oligomer sizes are presented as violin plots showing the kernel probability distribution, median and interquartile range for each treatment. The plots were generated, and statistical analysis was performed, using Prism8 (GraphPad, CA, USA). Data was analysed via an ANOVA with subsequent Kruskal-Wallis tests followed by Dunn’s multiple comparisons (P values are given). Stoichiometries of complexes were calculated by pairing of colocalised FPP for AF647-CLIC1_C24_ and AF488-αBc_C176_. Heatmaps were generated in MATLAB using home-written scripts.

## Results

### αBc binds and inhibits the amorphous aggregation of CLIC1 in vitro

We first assessed the ability of the sHsp αBc to prevent the heat-induced amorphous aggregation of a model client protein, CLIC1_C24_. When CLIC1_C24_ was incubated at 37°C, there was a significant increase in light scattering at 340 nm over 20 hr, indicative of protein aggregation (Fig. 1A). However, when CLIC1_C24_ was incubated in the presence of αBc_WT_ there was a concentration-dependent reduction in the rate and overall amount of light scatter associated with CLIC1_C24_ aggregation (Fig. 1A, B). The specificity of this effect was demonstrated by a negative control (using the non-chaperone protein SOD1) not impacting on the increase in light scatter associated with the aggregation of CLIC1_C24_ when incubated together. Furthermore, there was no increase in light scattering when αBc_WT_ or SOD1 were incubated alone, demonstrating that the increase in light scatter was exclusively due to the aggregation of CLIC1_C24_.

**Figure 1:**
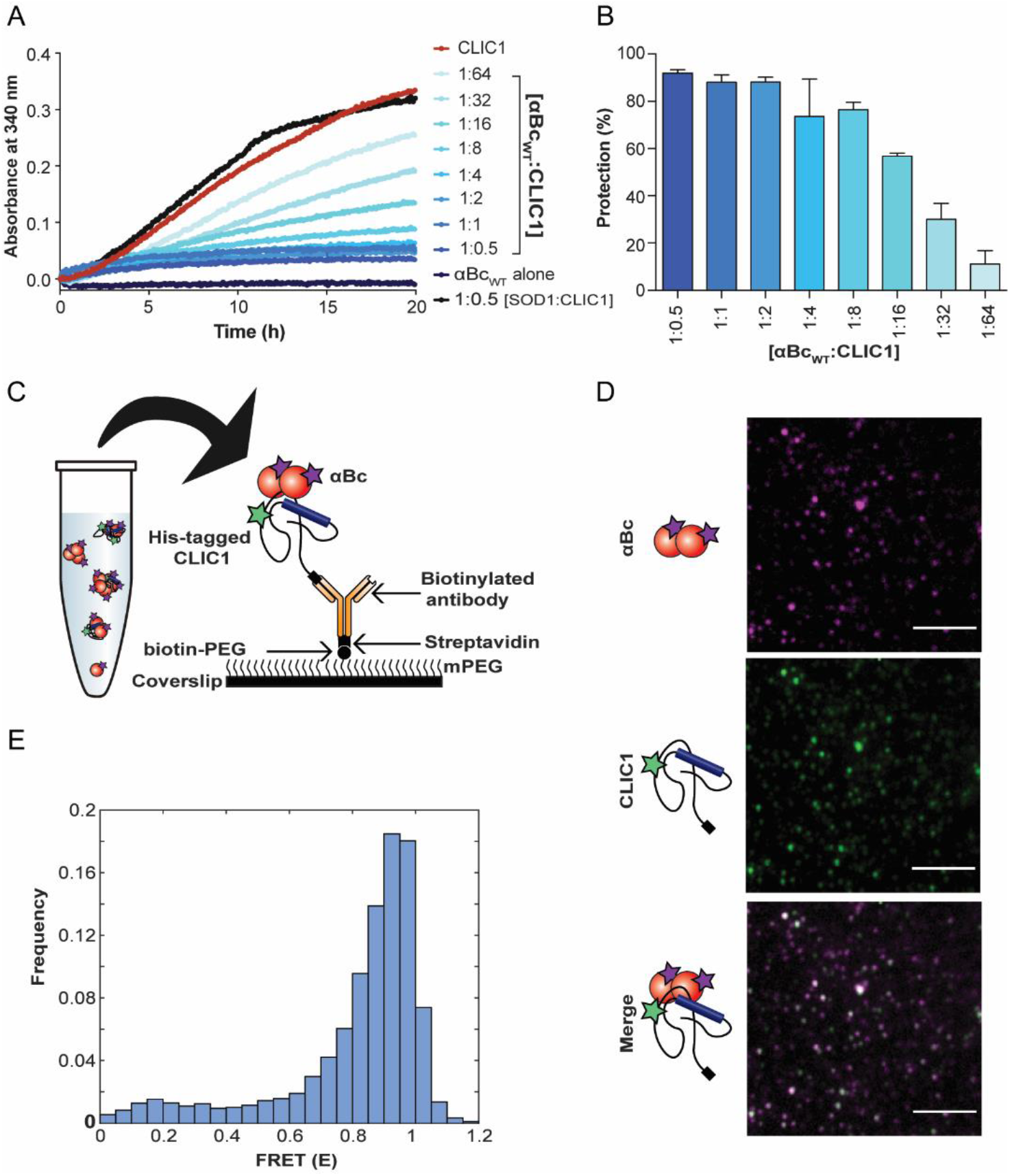
αBc binds and inhibits the amorphous aggregation of CLIC1 by forming stable client-chaperone complexes. **(A)** Recombinant CLIC1_C24_ was incubated at 37°C for 20 hr in the presence or absence of varying molar ratios of αBc_WT_ (1:0.5 to 1:64, αBc_WT_:CLIC1_C24_) or the control protein SOD1. The aggregation of CLIC1_C24_ was monitored by measuring the change in light scatter at 340 nm over time. **(B)** The percent inhibition afforded by varying molar ratios of αBc_WT_ against CLIC1 aggregation, reported as mean ± standard deviation of three independent experiments (n = 3). **(C)** Schematic of complex formation and surface immobilisation of complexes formed between AF555-CLIC1_C24_ and AF647-αBc_C176_ for smFRET experiments. **(D)** Representative z-stack TIRF microscopy images of AF555-CLIC1_C24_ and AF647-αBc_C176_ complexes. *Scale bar* 5 μim.**(E)** FRET efficiency *(E)* histogram derived from TIRF microscopy data of the initial intensities of CLIC1_C24_:αBc_C176_ complexes prior to photobleaching (n = 421 molecules).

To determine the nature of the physical interaction between CLIC1_C24_ and αBc_WT_, as suggested by the light scattering experiments, we utilised a single-molecule FRET based approach that allows interactions between biomolecules to be observed (at separations of 2-10 nm). In these experiments we used a mutant of αBc (αBc_C176_), that contained an additional cysteine for site-specific attachment of an Alexa Fluor 647 acceptor fluorophore. The addition of the C-terminal cysteine did not affect the ability of the chaperone to inhibit CLIC1_C24_ aggregation (Supplementary Fig. 1), and mass photometry measurements revealed there to be only a small shift in the oligomeric distribution of dye-labelled αBc_C176_ towards the formation of smaller species (Supplementary Fig. 2). To determine if fluorescently labelled αBc_C176_ could form client-chaperone complexes with CLIC1_C24_, donor (AF555) labelled CLIC1_C24_ and acceptor (AF647) labelled αBc_C176_ were incubated together at 37°C for 20 hr and immobilised on a functionalised coverslip for TIRF microscopy (Fig. 1C). Complexes containing co-localised CLIC1_C24_ and αBc_C176_ were observed at the single-molecule level (Fig. 1D) and the approximate time-FRET traces were calculated using the donor and acceptor fluorescence time-intensity traces (Supplementary Fig. 3A). The time-FRET trajectories initially displayed high FRET efficiencies, which gradually decreased over time, likely due to the photobleaching of multiple acceptor fluorophores within the αBc_C176_-CLIC1_C24_ complexes (Supplementary Fig. 3B). Analysis of the initial FRET efficiency of αBc_C176_-CLIC1_C24_ complexes prior to photobleaching showed these complexes had a high FRET efficiency (E = 0.8 – 1) and therefore were in close proximity, consistent with a stable interaction between αBc_C176_ and heat-destabilised CLIC1_C24_ (Fig. 1E). However, the complexity of these smFRET traces, as a result of multiple donor and acceptor fluorophores within the complexes means calculation of accurate distances between acceptor and donor fluorophores and the precise stoichiometries of αBc_C176_ and CLIC1_C24_ cannot readily be determined using this approach.

### A single-molecule fluorescence-based approach can be used to examine interactions between αBc and CLIC1

We sought to develop a single-molecule fluorescence-based assay that would enable the stoichiometries of αBc_C176_ and CLIC1_C24_ within complexes to be interrogated. To do so, we first incubated the site-specific fluorescently labelled CLIC1_C24_ (AF647-CLIC1_C24_) and αBc_C176_ (AF488-αBc_C176_) together at 37°C and collected aliquots at various time-points over a 10 hr period. Samples were then diluted and immediately immobilised to the coverslip surface (via the His-tag on CLIC1_C24_) for imaging using TIRF microscopy. As expected, αBc_C176_ (green) was observed to colocalise with CLIC1_C24_ molecules (purple) (Fig. 2A), indicative of the formation of stable complexes between these two proteins as observed in the smFRET experiments (Fig. 1D). The proportion of CLIC1_C24_ molecules colocalised with αBc_C176_ increased rapidly over 1 hr (Fig. 2B). Interestingly, after 4 hr the proportion of CLIC1_C24_ colocalised with αBc_C176_ reached a maximum of approximately 50%, demonstrating that not all CLIC1_C24_ molecules were in complex with αBc_C176_ under these experimental conditions.

**Figure 2:**
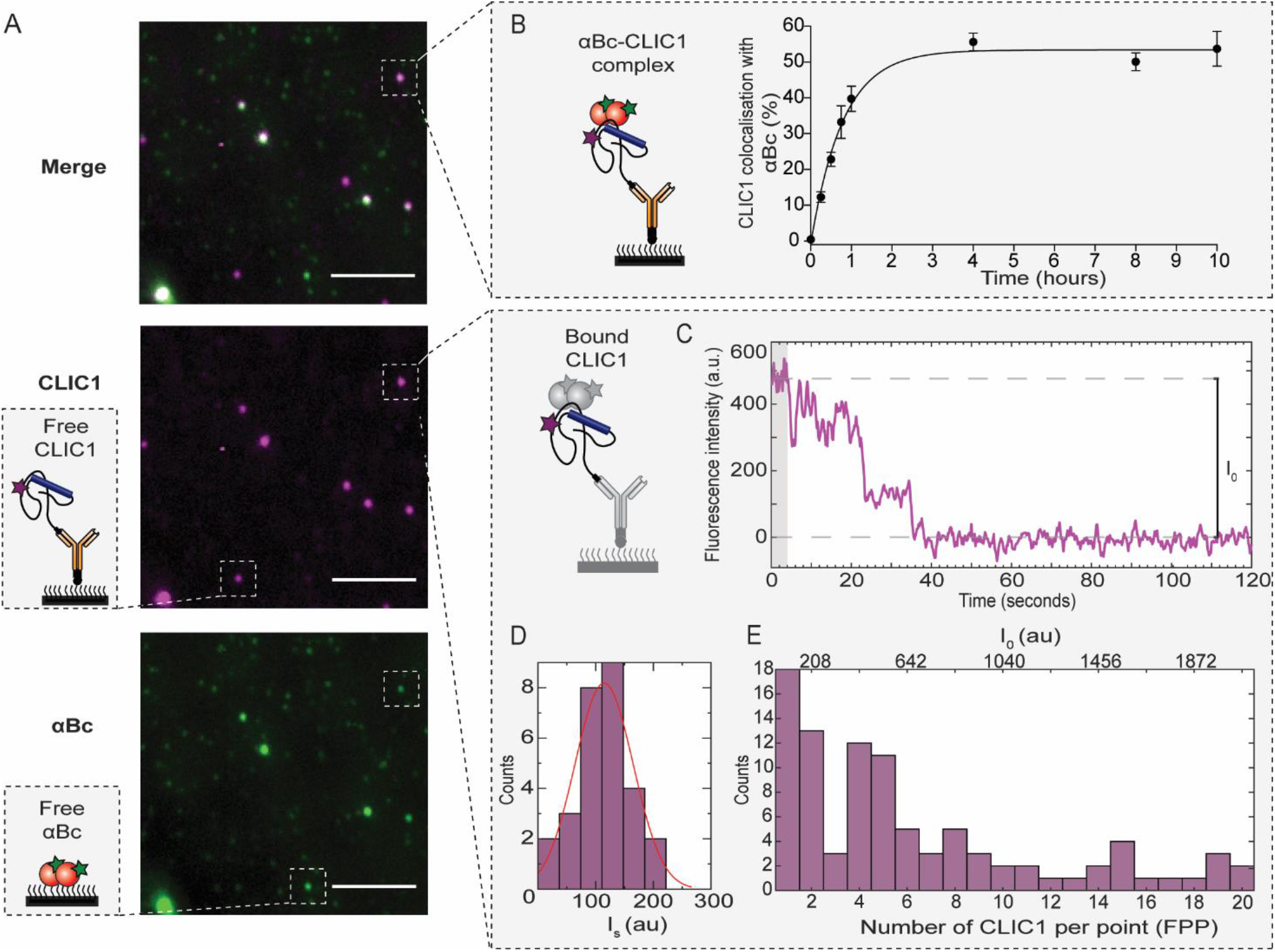
Characterisation of CLIC1_C24_-αBc_C176_ complexes using a single-molecule fluorescence-based approach. AF488-αBc_C176_ was incubated with AF647-CLIC1_C24_ (2:1 molar ratio) at 37°c for 10 hr to form complexes. Aliquots were taken at multiple timepoints throughout the incubation for TIRF microscopy imaging. **(A)** Representative TIRF microscopy images of complexes at 10 hr. *Scale bar* = 5 μm. *Schematic* indicating free CLIC1_C24_ and αBc_C176_ bound to the coverslip surface. **(B)** Schematic showing the immobilisation of αBc_C176_-CLIC1_C24_ complexes to the surface of a glass coverslip. The percentage of CLIC1_C24_ colocalised with αBc_C176_ over time reported as the mean ± standard error of the mean of three independent experiments. Data was fit using a one phase association model. **(C)** Example time trace of the fluorescent intensity of AF647-CLIC1_C24_ in complex with AF488-αBc_C176_. The shaded area (*grey*) represents the first 50 values that were averaged to determine the initial intensity (*I_0_*). **(D)** Photobleaching traces from individual AF647-CLIC1_C24_ molecules were manually identified and the I_s_ of each molecule was plotted and fit to a Gaussian distribution to determine the mean intensity of a single-photobleaching step (*I_s-mean_*). **(E)** Example histogram of CLIC1_C24_ showing the distribution of I_0_ and fluorescently-labelled proteins per point (FPP) at 10 h. FPP was calculated using the equation *FPP* = *I_0_*/*I_s-mean_* for all the CLIC1_C24_ in complex with αBc_C176_.

To determine the stoichiometries of CLIC1_C24_ and αBc_C176_ in these complexes, molecules were imaged until all fluorophores were completely photobleached, and the initial fluorescence intensity (*I_0_*) for both CLIC1_C24_ and αBc_C176_ in each complex determined (Fig. 2c, Supplementary Fig. 4, 5B). Individual monomers of CLIC1_C24_ and αBc_C176_ were identified manually by the presence of a distinct single photobleaching step and used to calculate the fluorescence intensity of a single-photobleaching event (*I_s_*) by fitting to a Gaussian distribution from which the mean (*I_s-mean_*) was derived (Fig. 2D, Supplementary Fig. 5c). The *I_s-mean_* were determined was determined to be 103 ± 52 a.u and 245 ± 62 a.u for CLIC1_C24_ and αBc_C176_ respectively, which were then used to determine the number of fluorescently-labelled proteins per point (FPP) for both CLIC1_C24_ and αBc_C176_. These FPP values were then used to determine the oligomer distribution of each protein at the time points examined (Fig. 2E, Supplementary Fig. 5D).

### The size and polydispersity of complexes formed between αBc and CLIC1 increase over time

To obtain further information on the interaction between αBc_C176_ and CLIC1_C24_, we examined the change in size and composition of the αBc_C176_-CLIC1_C24_ complexes over time, as well as the state of the molecules that were not in complex. Prior to incubation, both CLIC1_C24_ and αBc_C176_ were present predominantly as small non-colocalised monomers and dimers (Fig. 3A, 3E). Following incubation at 37°C for 0.25 hr, αBc_C176_ was found bound to oligomeric species of CLIC1_C24_ that were significantly larger in size compared to free CLIC1_C24_ species (Fig. 3B, P < 0.0001). The bound CLIC1_C24_ oligomers did not increase in size over the 10 hr incubation period (Fig. 3C). Conversely, the free CLIC1_C24_ molecules increased in size over time (P < 0.004) such that, after 10 hr they were of a similar size to the CLIC1_C24_ in complex with αBc_C176_, suggesting that this free CLIC1_C24_ aggregates to some extent during the incubation (Fig. 3D).

**Figure 3:**
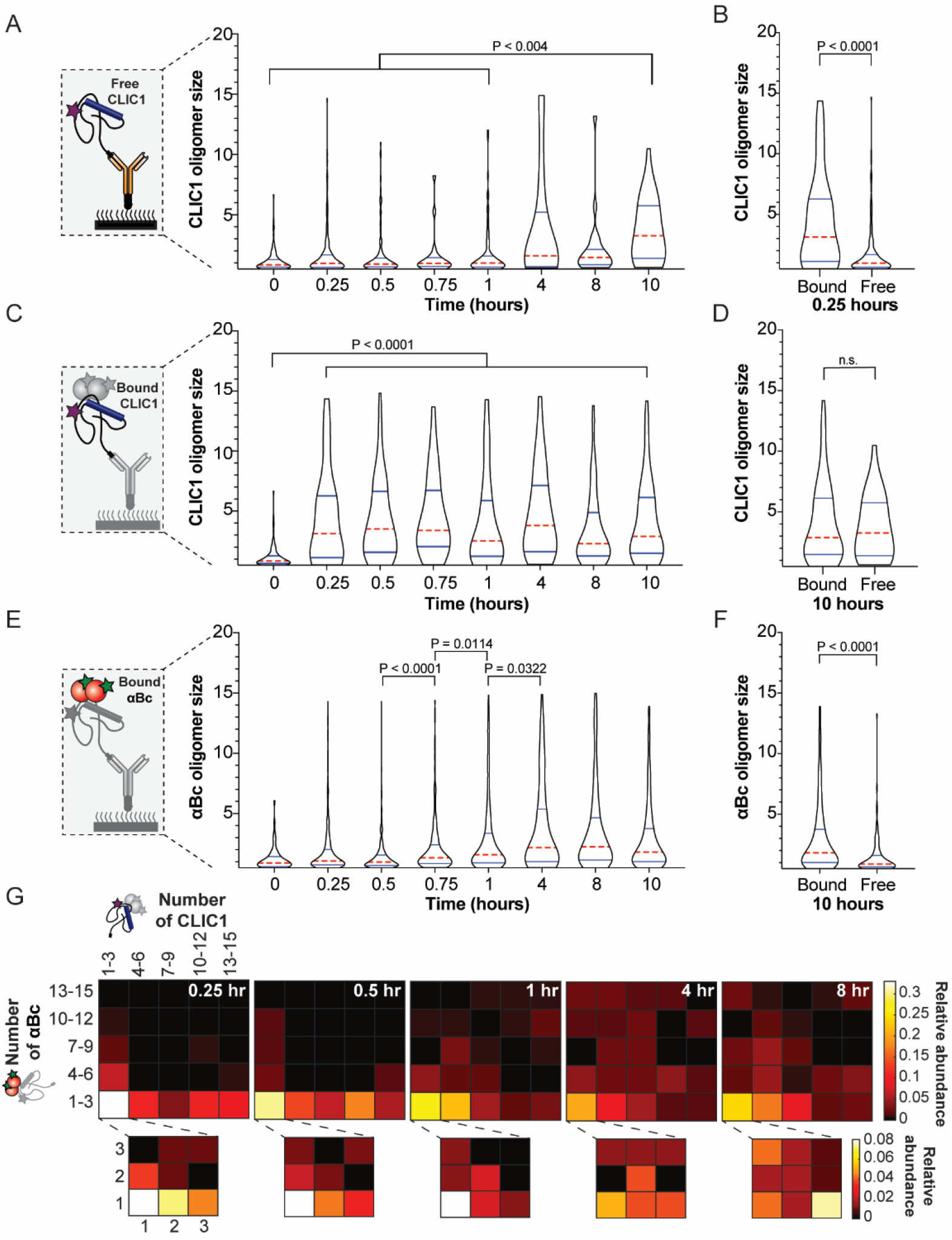
αBc_C176_-CLIC1_C24_ complexes increase in polydispersity and size over time. Violin plots showing the size distribution over 10 hr of **(A)** non-colocalised CLIC1_C24_ that is not in complex with αBc_C176_, **(B)** CLIC1_C24_ bound to αBc_C176_ or non-colocalised CLIC1_C24_ (Free) after 0.25 hr incubation, **(C)** CLIC1_C24_ bound to αBc_C176_, **(D)** CLIC1_C24_ bound to αBc_C176_ or non-colocalised CLIC1_C24_ (Free) after 10 hr incubation, **(E)** αBc_C176_ bound to CLIC1_C24_, and **(F)** αBc_C176_ bound to CLIC1_C24_ or non-specifically adsorbed to the surface (Free) after 10 hr. The violin plots show the kernel probability density (*black outline*), median (*red*) and interquartile range (*blue*). Results include measurements from three independent experiments (n = 3) and comparisons of distributions was performed using Kruskal-Wallis test for multiple comparisons with Dunn’s procedure (P values indicated). **(G)** Heat-maps showing the relative abundance of αBc_C176_-CLIC1_C24_ complexes and their stoichiometries over 8 hr of incubation.

During the early stages of the incubation (up to 0.5 hr), αBc_C176_ in complex with CLIC1_C24_ was primarily monomeric or dimeric (Fig. 3E). However, after 0.5 hr incubation the number of αBc_C176_ molecules in these complexes significantly increased over time, reaching a maximum after 4 hr. Despite having blocked (passivated) the coverslip surface, which significantly reduced the non-specific binding of αBc_C176_ to the coverslip, some non-specific binding was still observed (Supplementary Fig. 6C). Analysis of these non-specifically adsorbed αBc_C176_ species indicated that they were significantly smaller in size compared to αBc_C176_ that was in complex with CLIC1_C24_ after 10 hr (P < 0.0001; Fig. 3F, Supplementary Fig. 6D).

We next utilised our single-molecule fluorescence-based approach to characterise the stoichiometries of αBc_C176_-CLIC1_C24_ in individual complexes and interrogate how these change as a function of incubation time. For each individually identified αBc_C176_-CLIC1_C24_ complex, we determined the αBc_C176_:CLIC1_C24_ stoichiometry by calculating the number of monomers of each protein present. This process allowed us to quantify the relative abundance of these stoichiometries over time. Interestingly, we observed that complexes became increasingly polydisperse over the observation time (Fig. 3G). At early timepoints during the incubation (0.25 – 0.5 hr), complexes were comprised predominantly of smaller species of αBc_C176_ (monomers-3mers) bound to a polydisperse range of CLIC1_C24_ oligomers (monomers to 15mers). More specifically, the most abundant complex observed was comprised of monomeric αBc_C176_ bound to a single subunit of CLIC1_C24_. The polydispersity of CLIC1_C24_ within complexes (monomers to 15mers) did not change greatly over 8 hr, however, the relative abundance of complexes with more αBc_C176_ (> 6mers) increased after 1 hr. This increase in the stoichiometry of αBc_C176_:CLIC1_C24_ was consistent with the observed increase in the size distribution of αBc_C176_ over time (Fig. 3C). Together, these results suggest smaller αBc_C176_ subunits initially bind to aggregation-prone CLIC1_C24_ to form chaperone-client complexes and, over time, more free αBc_C176_ subunits bind to these complexes until the system reaches an equilibrium.

### Chaperone concentration influences the stoichiometries of CLIC1-αBc complexes

The molar ratio of sHsp to client protein is thought to be one of the most important parameters that determines the nature and size of sHsp-client complexes^14,22–24,26,39,40^. Therefore, we exploited our single-molecule fluorescence assay to investigate how sHsp concentration affects the stoichiometries of complexes formed with CLIC1_C24_. We observed that αBc_C176_ formed complexes with a range of CLIC1_C24_ oligomers and that there was no significant difference in the number of CLIC1_C24_ molecules in these complexes between the molar ratios tested (Fig. 4A). However, the number of αBc_C176_ subunits in these complexes was significantly smaller when the sHsp was present at lower molar ratios (0.5:1 or 1:1, αBc_C176_:CLIC1_C24_) compared to those formed at higher molar ratios (2:1 and 4:1, αBc_C176_: CLIC1_C24_) (P = 0.0024) (Fig. 4B). At all molar ratios tested, both αBc_C176_ and CLIC1_C24_ were significantly larger when in complex together compared to when freely bound to the surface (Supplementary Fig. 7). Interestingly, non-colocalised αBc_C176_ was observed to be significantly larger in size when incubated at the higher concentrations (4 μM) used in the molar-ratio of 4:1 (αBc_C176_: CLIC1_C24_) compared to lower concentrations (P = 0.0371) (Supplementary Fig. 7c).

**Figure 4:**
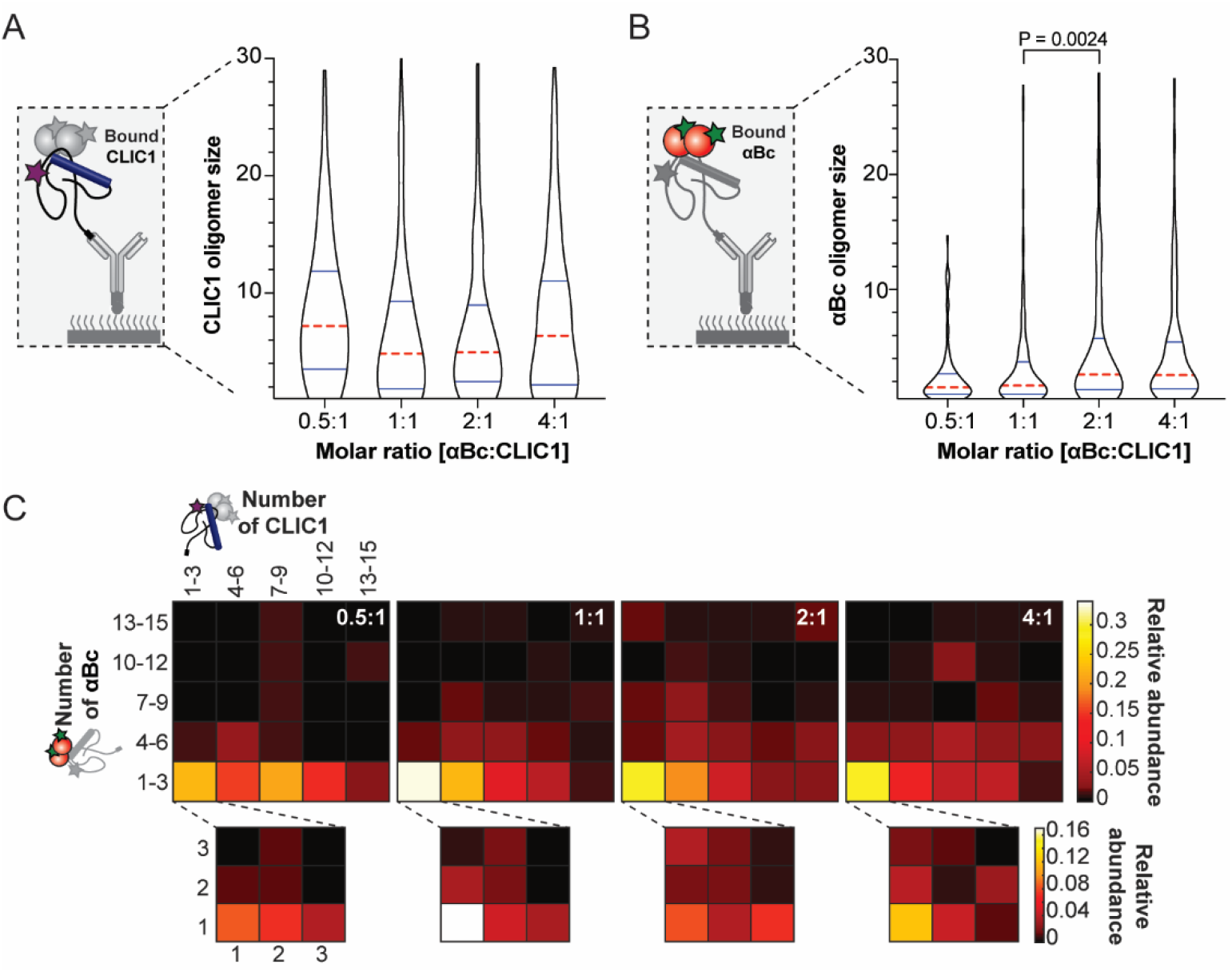
αBc_C176_-CLIC1_C24_ complexes change in size and stoichiometry with increasing αBc_C176_ concentration. The size distributions of CLIC1_C24_ **(A)** in complex with αBc_C176_ **(B)** at increasing molar ratios of αBc_C176_:CLIC1_C24_. The violin plots show the kernel probability density (*black outline*), median (*red*) and interquartile range shown (*blue*). Result are representative of three independent experiments (n = 3) and comparisons of distributions was performed using Kruskal-Wallis test for multiple comparisons with Dunn’s procedure (P values indicated). **(C)** Heat-maps showing the relative abundance of αBc_C176_-CLIC1_C24_ complexes with increasing molar ratios of αBc_C176_:CLIC1_C24_.

As observed previously, the complexes formed between αBc_C176_ and CLIC1_C24_ after heating were heterogeneous (Fig. 4C). Examination of the relative abundance of complexes formed when the molar ratio of αBc_C176_:CLIC1_C24_ was low ([0.5:1]-[1:1]) indicated that a small number of αBc_C176_ subunits (monomers-6mers) were in complex with a polydisperse range of CLIC1_C24_ species. In contrast, when complexes were formed at higher molar ratios of αBc_C176_:CLIC1_C24_ ([2:1]-[4:1]) more complexes contained a larger number of αBc_C176_ subunits (> 10mers). Consequently, this data suggests that higher concentrations of αBc_C176_ results in an increased binding of αBc_C176_ subunits to the complexes that are formed with CLIC1_C24_.

## Discussion

In this study we set out to detect and quantify for the first time the initial binding events between a sHsp and client protein. To do so we developed a single-molecule fluorescence assay to study the chaperone action of αBc, an archetypal mammalian sHsp. Employing such a single-molecule fluorescence-based approach we have determined the stoichiometries of complexes formed between αBc and a client protein, CLIC1. By examining the polydispersity and stoichiometries of these complexes over time, we have uncovered unique and important insights into the mechanism by which αBc captures misfolded client proteins to prevent their aggregation.

The most commonly used approach to investigate chaperone activity are assays that monitor the aggregation of proteins *in vitro*, either via light scatter or, in the case of amyloid fibril formation, fluorescent dyes such as Thioflavin T^41^. Indeed, we demonstrate via light scattering assay that αBc is able to effectively inhibit the heat-induced aggregation of CLIC1. However, these bulk ensemble assays struggle to provide mechanistic details concerning the interactions that occur between the chaperone and client protein which result in the suppression of aggregation. Furthermore, approaches such as size exclusion chromatography, electron microscopy and native mass spectrometry have traditionally been used to examine the end-stage complexes formed between sHsps and client proteins. These approaches are limited in their ability to capture the initial binding events between sHsps and client proteins and the dynamic nature of these complexes. In order to overcome these limitations, we developed a single-molecule fluorescence-based approach that, by utilising a step-wise photobleaching method, enables the stoichiometries of the chaperone-client complexes in solution to be revealed. In the case of αBc and CLIC1, by monitoring complexes in solution through time we have been able to uncover novel details of how this sHsp forms high-molecular mass complexes with client proteins.

We demonstrate that initially smaller species of αBc (predominantly monomers and dimers) bind to heat-destabilised CLIC1 oligomers. This observation validates previous suggestions, based on studying end-stage complexes, that smaller species of sHsps have high chaperone ability and therefore initially bind to misfolded proteins^18,42,43^. Interestingly, we observed that the number of complexes formed between αBc and CLIC1 increased rapidly over the first hour of incubation and reached a plateau after 4 hr. During this period there was an increase in the number of αBc subunits in each αBc-CLIC1 complex. We rationalise this as the recruitment of free αBc subunits onto existing αBc-CLIC1 complexes over time, as has been suggested to occur for other sHsp-client protein interactions^24,40,44^. Varying the molar ratio between CLIC1 and αBc, such that more αBc subunits were available to bind to CLIC1, resulted in an increase in size of these complexes. This time- and concentration-dependent recruitment of free αBc subunits onto αBc-CLIC1 complexes suggests that αBc is in an equilibrium between species bound to complexes and a constant pool of free subunits in solution.

It is known that many sHsps, including αBc, maintain large oligomeric assemblies via dynamic subunit exchange^8,45,46^. Moreover, in both prokaryotic (IbpA and IbpB)^44^ and eukaryotic sHsp systems (Hsp18.1 and Hsp16.6)^26^, sHsp-client complexes are dynamic in that sHsp subunits associate and dissociate from these complexes. Therefore, we propose that the observed accumulation of αBc onto αBc-CLIC1 complexes is regulated by the association and dissociation rates of the αBc subunits. Whilst we did not specifically probe for these dynamics in this study, the ability of single-molecule fluorescence techniques to observe dynamic and transient interactions in real-time provides the potential to further develop the approaches we have described here in order to examine if dynamic sHsp subunit exchange occurs on sHsp-client protein complexes.

Taken together, our findings provide direct experimental evidence for a two-step mechanism of sHsp-client complex formation that is in accordance with current models of sHsp chaperone action (Fig. 5)^24,47–49^. First, smaller (dissociated) sHsp species recognise and bind to misfolded client proteins. This allows for the subsequent addition of free sHsp subunits onto the newly formed complex until such a time that the system reaches an equilibrium between bound and unbound sHsps and no further growth of the complexes occurs. A two-step mechanism of chaperone action is also consistent with data obtained for plant sHsps and therefore may be a universal functional mechanism of sHsps that are able to form large oligomeric ensembles^24^. Future studies employing similar single-molecule fluorescence-based approaches to study the chaperone action of other polydisperse sHsps, such as Hsp27, will provide further insight into if this is indeed the case. Furthermore, similar studies that employ different client proteins would reveal whether the model of sHsp function described in this work is a general mechanism of sHsp/client interactions. Determining the precise molecular mechanisms of sHsps action is crucial to understanding how these molecular chaperones function to protect the cell from protein misfolding and their overall role in the cellular proteostasis network.

**Figure 5:**
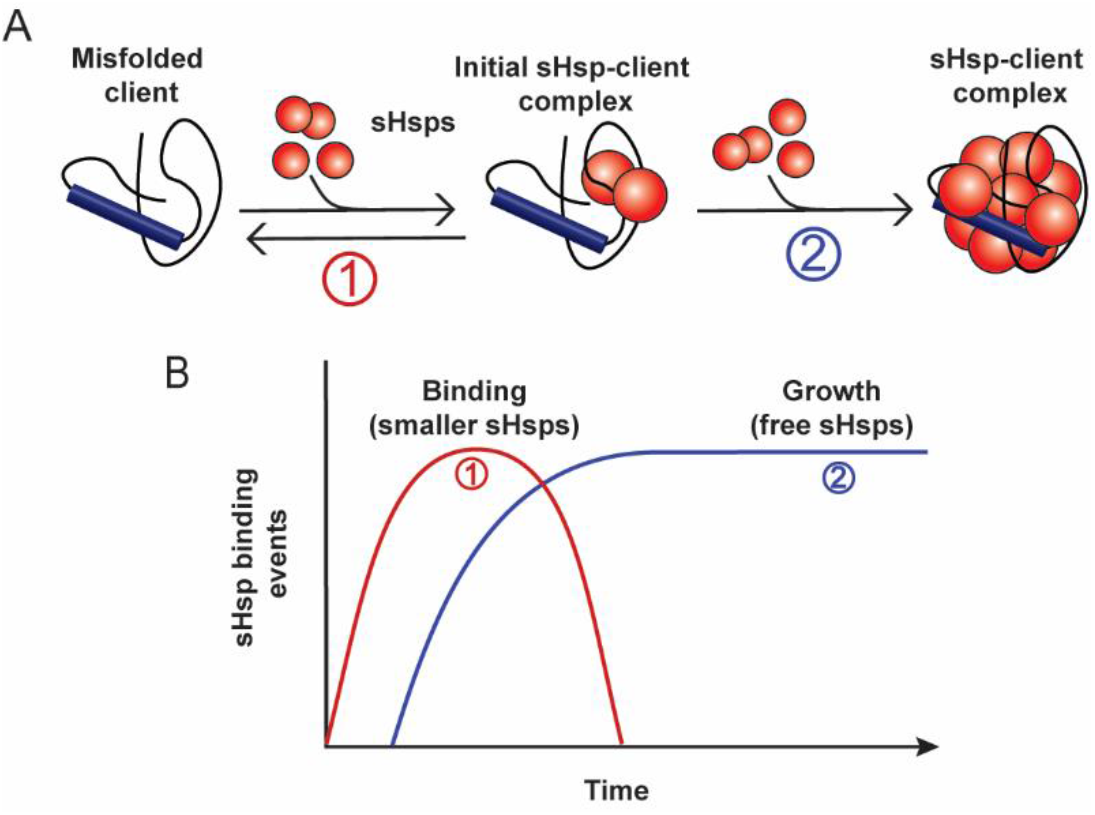
Schematic of two-step mechanism of sHsp-client complex formation. **(A)** Smaller sHsps initially recognise and bind misfolded client proteins (1) allowing for subsequent binding of additional sHsps subunits to form a large sHsp-client complex (2). **(B)** Theoretical binding events of sHsp subunits over time showing that initial binding of sHsps to their clients increases over time (1) until all the misfolded client is bound and additional sHsp subunits associate with these complexes (2) in order to form sHsp-client complexes.

## Author contributions

CLJ, HE and AVO formulated the experimental approach. NM made the recombinant CLIC1 and performed *In vitro* aggregation assays. GW and JB performed mass photometry experiments. CLJ performed all other experiments, analysed the data, constructed the figures and wrote the initial manuscript. CLJ, NM, BP, JB, HE and AVO edited the manuscript and approved the submission of the final manuscript.

## Acknowledgements

This research performed by CJ has been conducted with the support of the Australian Government Research Training Program Scholarship. We thank Drs Philipp Kukura and Weston Struwe (University of Oxford, UK) for their help with the mass photometry. We also thank Dr Sophie Goodchild (Macquarie University, Australia) and Prof Paul Curmi (UNSW, Australia) for their help and guidance with the design of the CLIC1 mutant proteins used in this work. Finally we thank the Illawarra Heath and Medical Research Institute for technical and administrative support.

## References

1. Jakob, U., Gaestel, M., Engel, K. & Buchner, J. Small heat shock proteins are molecular chaperones. Journal of Biological Chemistry 268, 1517–1520 (1993).

2. Horwitz, J. Alpha-crystallin can function as a molecular chaperone. P Natl A Sci 89, 10449–10453 (1992).

3. de Jong, W.W., Leunissen, J.A. & Voorter, C. Evolution of the alpha-crystallin/small heat-shock protein family. Molecular biology and evolution 10, 103–126 (1993).

4. Clark, J.I. & Muchowski, P.J. Small heat-shock proteins and their potential role in human disease. Current opinion in structural biology 10, 52–59 (2000).

5. Ciocca, D.R. & Calderwood, S.K. Heat shock proteins in cancer: diagnostic, prognostic, predictive, and treatment implications. Cell stress & chaperones 10, 86 (2005).

6. Sun, Y. & MacRae, T.H. The small heat shock proteins and their role in human disease. The FEBS journal 272, 2613–2627 (2005).

7. Basha, E., O’Neill, H. & Vierling, E. Small heat shock proteins and α-crystallins: dynamic proteins with flexible functions. Trends in Biochemical Sciences 37, 106–117 (2012).

8. Aquilina, J.A., Benesch, J.L.P., Bateman, O.A., Slingsby, C. & Robinson, C.V. Polydispersity of a mammalian chaperone: mass spectrometry reveals the population of oligomers in αB-crystallin. P Natl A Sci 100, 10611–10616 (2003).

9. Baldwin, A.J., Lioe, H., Robinson, C.V., Kay, L.E. & Benesch, J.L. αB-crystallin polydispersity is a consequence of unbiased quaternary dynamics. Journal of molecular biology 413, 297–309 (2011).

10. Haslbeck, M., Weinkauf, S. & Buchner, J. Regulation of the chaperone function of small Hsps. in The big book on small heat shock proteins 155–178 (Springer, 2015).

11. Jovcevski, B. et al. Phosphomimics destabilize Hsp27 oligomeric assemblies and enhance chaperone activity. Chem Biol 22, 186–195 (2015).

12. Lindner, R.A., Kapur, A. & Carver, J.A. The interaction of the molecular chaperone, α-crystallin, with molten globule states of bovine α-lactalbumin. Journal of Biological Chemistry 272, 27722–27729 (1997).

13. Regini, J.W. et al. The interaction of unfolding α-lactalbumin and malate dehydrogenase with the molecular chaperone αB-crystallin: a light and X-ray scattering investigation. Molecular Vision 16, 2446–2456 (2010).

14. Stromer, T., Ehrnsperger, M., Gaestel, M. & Buchner, J. Analysis of the interaction of small heat shock proteins with unfolding proteins. Journal of Biological Chemistry 278, 18015–18021 (2003).

15. Giese, K.C. & Vierling, E. Changes in oligomerization are essential for the chaperone activity of a small heat shock protein in vivo and in vitro. The Journal of Biological Chemistry 277, 46310–46318 (2002).

16. Shashidharamurthy, R., Koteiche, H.A., Dong, J. & McHaourab, H.S. Mechanism of chaperone function in small heat shock proteins: dissociation of the HSP27 oligomer is required for recognition and binding of destabilized T4 lysozyme. Biol Chem 280, 5281–5289 (2005).

17. McHaourab, H.S., Godar, J.A. & Stewart, P.L. Structure and mechanism of protein stability sensors: chaperone activity of small heat shock proteins. Biochemistry 48, 3828–3837 (2009).

18. Santhanagopalan, I. et al. It takes a dimer to tango: Oligomeric small heat shock proteins dissociate to capture substrate. Journal of Biological Chemistry 293, 19511–19521 (2018).

19. Aquilina, J.A. et al. Subunit exchange of polydisperse proteins: mass spectrometry reveals consequences of alphaA-crystallin truncation. Journal of Biological Chemistry 280, 14485–11491 (2005).

20. Augusteyn, R.C. Dissociation is not required for alpha-crystallin’s chaperone function. Experimental Eye Research 79, 781–784 (2004).

21. Franzmann, T., Wühr, M., Richter, K., Walter, S. & Buchner, J. The activation mechanism of Hsp26 involves structural changes within the oligomer. Dissociation is not required. J. Mol. Biol 350, 1083–1093 (2005).

22. Basha, E., Lee, G.J., Demeler, B. & Vierling, E. Chaperone activity of cytosolic small heat shock proteins from wheat. European Journal of Biochemistry 271, 1426–1436 (2004).

23. Lee, G.J., Roseman, A.M., Saibil, H.R. & Vierling, E. A small heat shock protein stably binds heat-denatured model substrates and can maintain a substrate in a folding-competent state. EMBO J. 16, 659–671 (1997).

24. Stengel, F. et al. Quaternary dynamics and plasticity underlie small heat shock protein chaperone function. Proceedings of the National Academy of Sciences of the United States of America 107, 2007–2012 (2010).

25. Stengel, F. et al. Dissecting heterogeneous molecular chaperone complexes using a mass spectrum deconvolution approach. Chemistry & biology 19, 599–607 (2012).

26. Friedrich, K.L., Giese, K.C., Buan, N.R. & Vierling, E. Interactions between small heat shock protein subunits and substrate in small heat shock protein-substrate complexes. J. Biol. Chem. 279, 1080–1089 (2004).

27. van Montfort, R.L., Basha, E., Friedrich, K.L., Slingsby, C. & Vierling, E. Crystal structure and assembly of a eukaryotic small heat shock protein. Nature Structural & Molecular Biology 8, 1025–1030 (2001).

28. Johnston, C.L., Marzano, N.R., van Oijen, A.M. & Ecroyd, H. Using single-molecule approaches to understand the molecular mechanisms of heat-shock protein chaperone function. Journal of molecular biology 430, 4525–4546 (2018).

29. Fanucchi, S., Adamson, R.J. & Dirr, H.W. Formation of an unfolding intermediate state of soluble chloride intracellular channel protein CLIC1 at acidic pH. Biochemistry 47, 11674–11681 (2008).

30. Cross, M., Fernandes, M., Dirr, H. & Fanucchi, S. Glutamate 85 and glutamate 228 contribute to the pH-response of the soluble form of chloride intracellular channel 1. Molecular and cellular biochemistry 398, 83–93 (2015).

31. Haslbeck, M. & Vierling, E. A first line of stress defense: small heat shock proteins and their function in protein homeostasis. Journal of molecular biology 427, 1537–1548 (2015).

32. Horwitz, J., Huang, Q.-L., Linlin, D. & Bova, M.P. Lens alpha-crystallin: chaperone-like properties. Method Enzymol 290, 365–383 (1998).

33. Kim, Y. et al. Efficient site-specific labeling of proteins via cysteines. Bioconjugate chemistry 19, 786–791 (2008).

34. Chandradoss, S.D. et al. Surface passivation for single-molecule protein studies. JoVE (Journal of Visualized Experiments), e50549 (2014).

35. Pan, H., Xia, Y., Qin, M., Cao, Y. & Wang, W. A simple procedure to improve the surface passivation for single molecule fluorescence studies. Physical biology 12, 045006 (2015).

36. Zhong, Y. et al. CHD4 slides nucleosomes by decoupling entry-and exit-side DNA translocation. bioRxiv, 684860 (2019).

37. Hadzic, M.C., Kowerko, D., Börner, R., Zelger-Paulus, S. & Sigel, R.K. Detailed analysis of complex single molecule FRET data with the software MASH. in Imaging, Manipulation, and Analysis of Biomolecules, Cells, and Tissues IX Vol. 9711 971119 (International Society for Optics and Photonics, 2016).

38. Schindelin, J. et al. Fiji: an open-source platform for biological-image analysis. Nature methods 9, 676 (2012).

39. Mogk, A. et al. Refolding of substrates bound to small Hsps relies on a disaggregation reaction mediated most efficiently by ClpB/DnaK. Journal of Biological Chemistry 278, 31033–31042 (2003).

40. Haslbeck, M. et al. Hsp26: a temperature-regulated chaperone. European Molecular Biology Organization Journal 18, 6744–6751 (1999).

41. Gade Malmos, K. et al. ThT 101: a primer on the use of thioflavin T to investigate amyloid formation. Amyloid 24, 1–16 (2017).

42. Hochberg, G.K. et al. The structured core domain of αB-crystallin can prevent amyloid fibrillation and associated toxicity. P Natl A Sci 111, E1562–E1570 (2014).

43. Alderson, T.R. et al. Local unfolding of the HSP27 monomer regulates chaperone activity. Nature Communications 10, 1068 (2019).

44. Żwirowski, S. et al. Hsp70 displaces small heat shock proteins from aggregates to initiate protein refolding. The EMBO journal 36, 783–796 (2017).

45. Van den Oetelaar, P.J., Van Someren, P.F., Thomson, J.A., Siezen, R.J. & Hoenders, H.J. A dynamic quaternary structure of bovine. alpha.-crystallin as indicated from intermolecular exchange of subunits. Biochemistry 29, 3488–3493 (1990).

46. Bova, M.P., Ding, L.L., Horwitz, J. & Fung, B.K. Subunit exchange of alphaA-crystallin. Biol Chem 272, 29511–29517 (1997).

47. Treweek, T.M., Meehan, S., Ecroyd, H. & Carver, J.A. Small heat-shock proteins: important players in regulating cellular proteostasis. Cell Mol Life Sci 72, 429–51 (2015).

48. Mogk, A., Ruger-Herreros, C. & Bukau, B. Cellular functions and mechanisms of action of small heat shock proteins. Annual review of microbiology 73 (2019).

49. Haslbeck, M., Weinkauf, S. & Buchner, J. Small heat shock proteins: Simplicity meets complexity. Journal of Biological Chemistry 294, 2121–2132 (2019).

