## Supplementary data for "Single-molecule fluorescence-based approach reveals novel mechanistic insights into small heat shock protein chaperone function"

### Supplementary Information and Figures

**Supplementary Table 1. Summary of labelling methods and efficiencies**

| **Protein** | **Label/dye** | **Method** | **Wavelength** | ***ε*** **(mg$^{-1}$ ml cm$^{-1}$)** | **Labelling efficiency** |
|---|---|---|---|---|---|
| CLIC1$_{C24}$ | Alexa Fluor 647-$C_2$-maleimide | UV abs | 650 nm | 0.55 | 96% |
| | Alexa Fluor 555-$C_5$-maleimide | UV abs | 556 nm | 0.55 | 82% |
| αBc$_{C176}$ | Alexa Fluor 647-$C_2$-maleimide | UV abs | 650 nm | 0.55 | 77% |
| | Alexa Fluor 488-$C_5$-maleimide | Mass spec | | | >95% |

* Method = method used to determine labelling efficiency (UV absorbance at wavelength specified or denatured mass spectrometry).

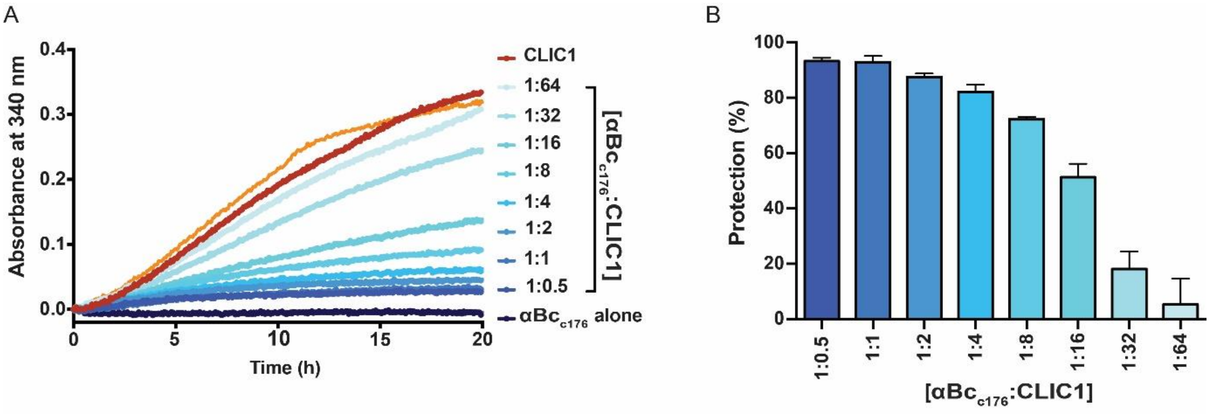


**Supplementary Figure 1: αBc$_{c176}$ inhibits the amorphous aggregation of CLIC1$_{C24}$. (A)** Recombinant CLIC1$_{C24}$ (30 μM) was incubated in 50 mM phosphate buffer (pH 7.4) at 37°C in the presence or absence of varying molar ratios of αBc$_{C176}$:CLIC1$_{C24}$ ranging from 1:0.5 to 1:64 for 20 hours. The CLIC1$_{C24}$ aggregation was monitored via the change in light scatter at 340 nm over time. **(B)** The percentage protection afforded by varying molar ratios of αBc$_{C176}$ against CLIC1 aggregation, reported as mean ± standard deviation of three independent experiments (n = 3).

*Mass photometry methodology*

The samples of αB-crystallin were diluted from stock to 2 μM (monomer), in 50 mM filtered phosphate buffer (pH 7.4). These solutions were incubated for 30 min at room temperature, then 45 min on ice. Further dilution to 400 nM (monomer) was performed immediately before measurement.

Coverslips were cleaned by sequential sonication in milli-Q water, isopropanol and milli-Q water (5 min each) before washing with ethanol and drying under a clean stream of nitrogen. Coverslips were then assembled into flow chambers as described previously (Young, Hundt et al. 2018).

Mass photometry measurements were made on a prototype of the ONE$^{MP}$ (Refeyn Ltd). For the mass photometry measurement, 10-15 mL of sample was added to the flow chamber and recording initiated as soon as the sample stage returned to the focal position. Two 60-sec recordings were taken for each sample, at an effective frame rate of 318 Hz and an effective pixel size of 21.1 nm. The resultant movies were analysed using software written in-house according to previously described procedures (Young, Hundt et al. 2018) using stacks of 15 frames to produce the averages before division.

To convert peak contrasts to mass, 15 standard proteins were measured on the same instrument under the same conditions, and the average contrast for each standard was plotted against the sequence mass to produce a linear calibration curve. The converted peak contrasts were then visualised in mass histograms (Supplementary Figure 2).

*References:*

Young, G., N. Hundt, D. Cole, A. Fineberg, J. Andrecka, A. Tyler, A. Olerinyova, A. Ansari, E. G. Marklund and M. P. Collier (2018). "Quantitative mass imaging of single biological macromolecules." Science **360**(6387): 423-427.

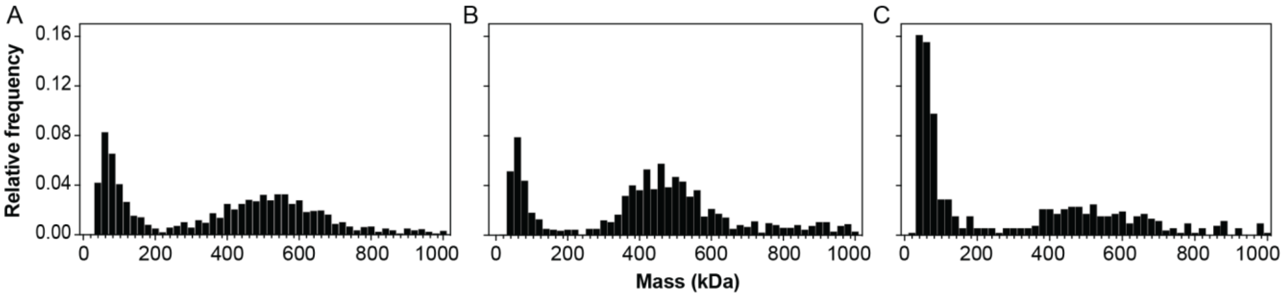


**Supplementary Figure 2: αBc variants size distributions determined by mass photometry.** Size distributions of **(A)** αBc$_{WT}$, **(B)** αBc$_{C176}$ and **(C)** Alexa Fluor 488-labelled αBc$_{C176}$. Oligomers of the different forms of αBc are seen in the range 300-800 kDa, with some sub-oligomeric species observed for all three proteins at 400 nM (30 min after dilution from 2 μM at room temperature). Fluorescent labelling leads to some dissociation of the large oligomers, but the majority of AF-488 αBc$_{C176}$ monomers remain in large oligomers.

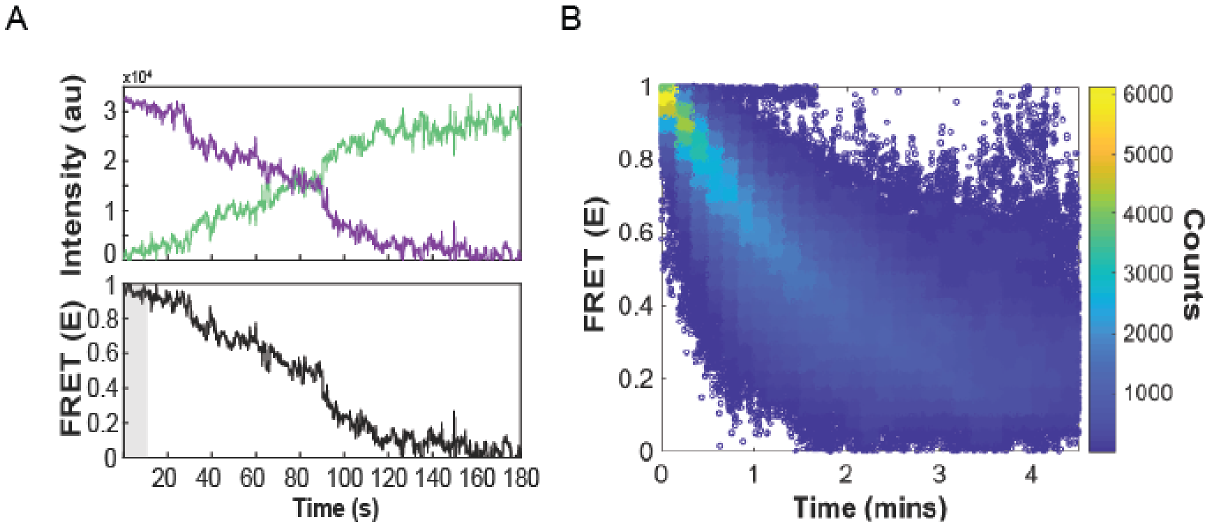


**Supplementary Figure 3: αBc$_{c176}$ and CLIC1$_{C24}$ form complexes that FRET. (A)** A representative smFRET trace of the fluorescence intensity of the donor AF555-CLIC1$_{C59S}$ (*green*) and acceptor AF647- αBc$_{C176}$ (*purple*) in complex over time. These intensity traces were used to calculate the FRET efficiency over time (*black*). *Grey area* represents the first 20 values that were used to construct FRET efficiency histogram for CLIC1$_{C24}$-αBc$_{C176}$ complexes. **(B)** FRET efficiency heatmap (n = 421 molecules) of CLIC1$_{C24}$-αBc$_{C176}$ complexes over time.

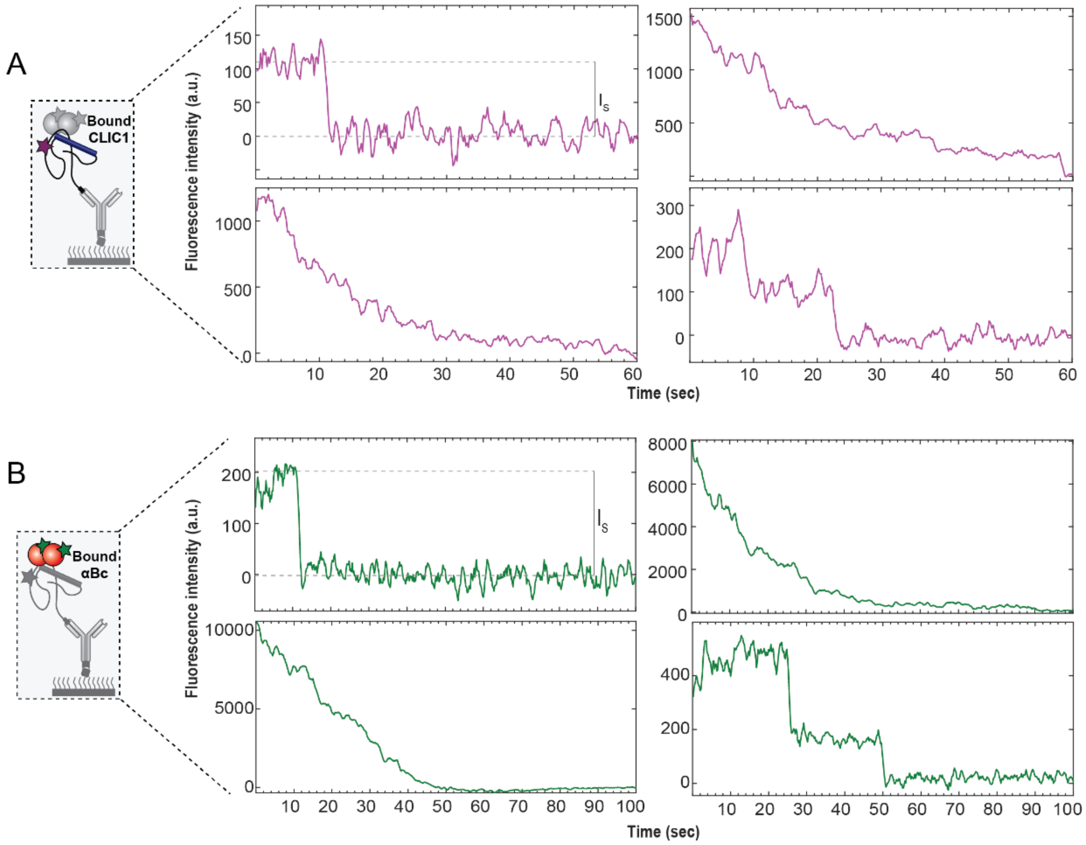


**Supplementary Figure 4:** Example time traces of the fluorescent intensity of multiple **(A)** Alexa Fluor 647-labelled CLIC1$_{C24}$ molecules or **(B)** Alexa Fluor 488-labelled αBc$_{C176}$ molecules in complex. Example of a trajectory with a single-photobleaching step ($I_s$) is shown for both CLIC1$_{C24}$ and αBc$_{c176}$.

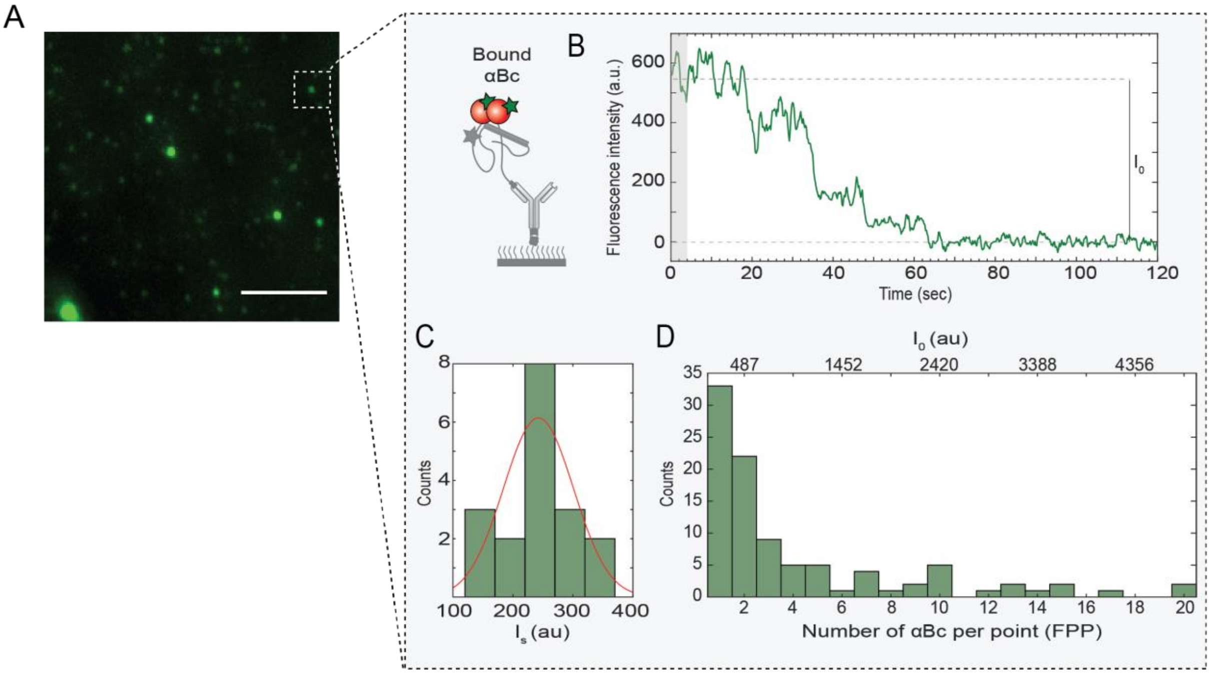


**Supplementary Figure 5: Determination of the size distribution of αBc$_{c176}$ in complex with CLIC1$_{C24}$ using our single-molecule fluorescence-based assay. (A)** Representative TIRF microscopy image of Alexa Fluor 488-labelled αBc$_{C176}$ in complex with CLIC1$_{C24}$ at 10 hr. *Scale bar* = 2 µm. **(B)** Example time trace of the fluorescence intensity of Alexa Fluor 488-labelled αBc$_{c176}$ in complex with CLIC1$_{C24}$. *Grey areas* represent the first 50 values averaged to determine the initial intensity. **(C)** αBc$_{C176}$ single-photobleaching intensity traces were manually identified and the initial intensities ($I_s$) of each was calculated and fit to a Gaussian distribution from which the mean ($I_{s\text{-mean}}$) was derived. **(D)** Example size distribution of showing the distribution of $I_0$ and number of αBc$_{C176}$ at 10 hr timepoint. The number of αBc$_{C176}$ was calculated by $I_0/I_{s\text{-mean}}$ for all the number of αBc$_{C176}$ in complex with αBc$_{C176}$ at 10 hr.

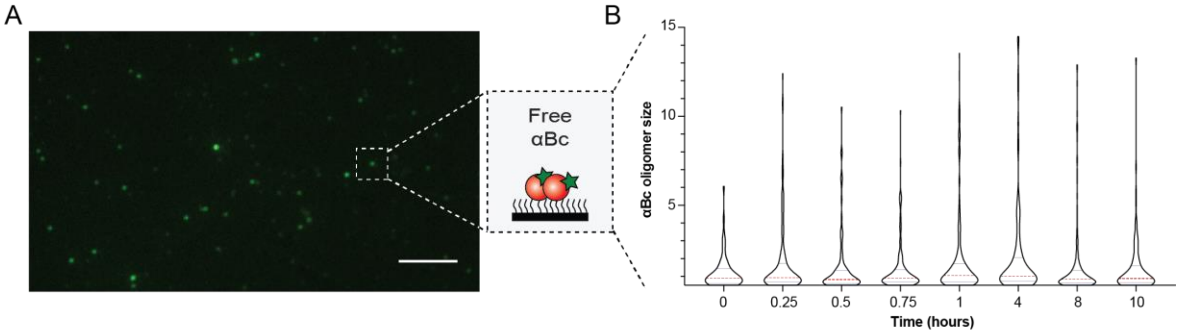


**Supplementary Figure 6: Non-colocalised αBc$_{c176}$ does not change in size over time. (A)** Example image of non-specific binding of Alexa Fluor 488-labelled αBc$_{C176}$ to blocked coverslip surface. Alexa Fluor 488-labelled αBc$_{C176}$ (20 nM) was incubated in flow cell for 5 min, washed with imaging buffer and imaged using TIRF microscopy. **(B)** Size distributions of non-specifically surface bound αBc$_{C176}$ at multiple timepoints over 10 hr. The violin plots show the kernel probability density (*black outline*), median (*red*) and interquartile range shown (*blue*). Result are representative of three independent experiments (n = 3).

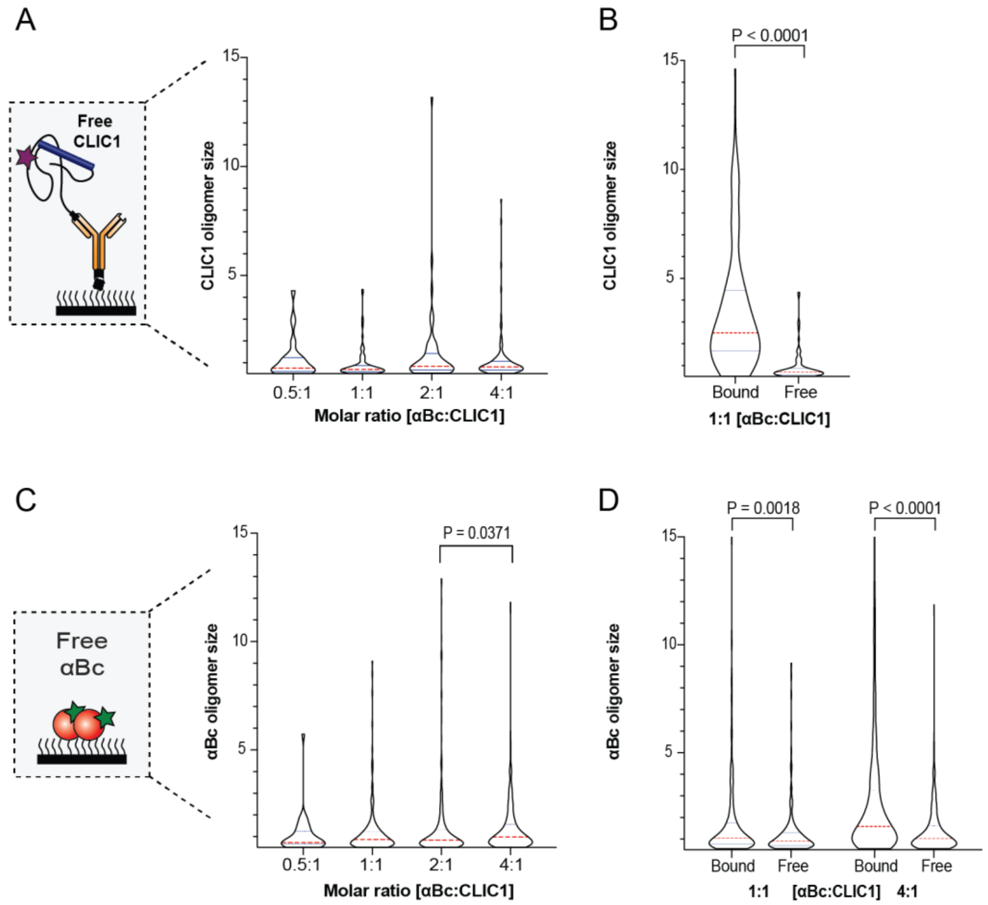


**Supplementary Figure 7: CLIC1$_{C24}$ and αBc$_{c176}$ are significantly smaller when not in complex compared to when they are in complex. (A)** Size distributions of free Alexa Fluor 647-labelled CLIC1$_{C24}$ not in complex with αBc$_{C176}$ at increasing molar ratios of αBc:CLIC1. **(B)** Size distributions of CLIC1$_{C24}$ bound to αBc$_{C176}$ compared to free CLIC1$_{C24}$ on the surface at 1:1 molar ratio. **(C)** Size distribution of non-specifically surface bound Alexa Fluor 488-labelled αBc$_{C176}$ at increasing molar ratios. **(D)** Comparison of size distributions of αBc$_{C176}$ bound to CLIC1$_{C24}$ and non-specifically bound (free) on the surface at 1:1 (*left*) and 4:1 (*right*) molar ratios [αBc$_{C176}$: CLIC1$_{C24}$]. The violin plots show the kernel probability density (*black outline*), median (*red*) and interquartile range shown (*blue*). Result are representative of three independent experiments (n = 3) and comparisons of distributions was performed using Kruskal-Wallis test for multiple comparisons with Dunn's procedure (P values indicated).
